## Supplementary materials for "Opponent intracerebral signals for reward and punishment prediction errors in humans"

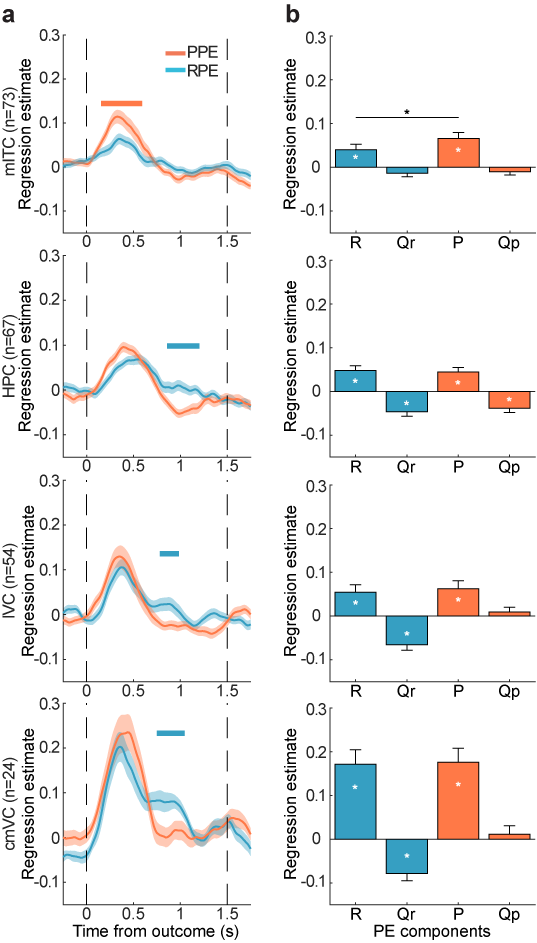

**Figue S1. Reward and punishment PE signals in supplementary PE regions. a.** Time course of regression estimates obtained from linear fit of BGA with PE modeled separately for the reward and punishment conditions. Horizontal bold lines indicate significant difference between conditions (blue: RPE>PPE; red: PPE>RPE; p_c_<0.05). Shaded areas represent inter-patient SEM. **b.** Regression estimates of broadband gamma power against prediction error components averaged over the 0.25-1 s post-outcome time window. lVC: lateral visual cortex; cmVC: caudal medial visual Cortex; mITC: medial inferior temporal cortex; HPC: hippocampus.

**Table S1 : Demographical and clinical details.**

Abbreviations used: orbitofrontal cortex: OFC; premotor cortex: PM; hippocampus: HPC; left: L; right: R; alprazolam: APZ; carbamazepine: CBZ; clobazam: CLB; gabapentin: GBP; lacosamide: LCM, lamotrigine: LMT; levetiracetam: LEV, nitrazepam: NZP; oxcarbazepine: OXC; perampanel: PER; topiramate: TPM; sodium valproate: VPA; zonisamide: ZON.

| **Patient** | **Sex** | **Age** | **Hand laterality** | **Number of iEEG sites** | **Epileptic focus** | **Epilepsy onset (age)** | **Antiepileptic drugs** |
| --- | --- | --- | --- | --- | --- | --- | --- |
| 1 | F | 29 | R | 98 | L frontopolar | 19 | LCM + LMT |
| 2 | F | 42 | R | 104 | L temporal | 13 | CLB + LEV + ZON |
| 3 | F | 20 | R | 98 | L temporal | 11 | CLB + LCM + LMT |
| 4 | M | 48 | L | 98 | R temporal | 12 | OXC |
| 5 | M | 37 | R | 100 | L OFC | 4 | LCM + NZP |
| 6 | F | 13 | R | 80 | R PM | 10 | CBZ + VPA |
| 7 | F | 34 | R | 80 | L HPC + insular | 5 | LCM + LEV |
| 8 | M | 32 | R | 96 | Bilateral temporal | 21 | LTG + PER |
| 9 | F | 15 | R | 91 | R temporal + insular | 7 | CLB + LEV |
| 10 | M | 28 | R | 155 | R fronto-temporal | 11 | CBZ + LCM |
| 11 | M | 20 | R | 161 | G frontal + temporal (2) | 14 | LCM + LMT + ZON |
| 12 | M | 41 | R | 135 | R temporal | 32 | CLB + GBP |
| 13 | F | 15 | R | 99 | L temporo-occipital | 12 | CLB + LCM + LMT |
| 14 | M | 40 | R | 110 | R temporal | 5 | CBZ + LCM |
| 15 | F | 47 | R | 101 | R insular + L frontal | 7 | LCM + TPM |
| 16 | F | 34 | R | 90 | R OFC + temporal | 28 | CBZ + LCM + LMT |
| 17 | M | 57 | R | 107 | L temporal | 37 | CBZ + LMT + PER |
| 18 | M | 41 | L | 45 | L temporal |  | CBZ + CLB |
| 19 | M | 29 | R | 99 | R OFC | 25 | LCM + LMT |
| 20 | F | 48 | R | 136 | R temporal | 19 | LCM + LEV + LMT |

**Table S2: Significance of prediction-error signals for the entire dataset.**

Areas (MarsAtlas labels) are ordered according to the t-value obtained by testing against zero (across sites within each parcel) the regression estimates of BGA against prediction errors (in the 0-1000 ms post-outcome time window). Bold p-values indicate significance after correction for multiple comparisons across regions (n=39 areas; α_corrected_=1.3×10^-3^ after Bonferroni correction). Grey: areas including either less than 9 recorded sites or a proportion of significant contact inferior to 0.2. lVC: lateral visual cortex; cmVC: caudal medial visual Cortex; mITC: medial inferior temporal cortex; HPC: hippocampus. pINS: posterior Insula.

| **Parcellation label** | **Number of iEEG sites** | **Number of patients** | **t-value** | **p-value** | **Number of significant patients** | **Proportion of significant contacts** |
| --- | --- | --- | --- | --- | --- | --- |
| **aINS** | 83 | 13 | 11,89 | **1,59×10^-19^** | 11 | 0,48 |
| **dlPFC** | 74 | 9 | 7,72 | **4,69×10^-11^** | 8 | 0,38 |
| **cmVC** | 24 | 7 | 6,65 | **8,81×10^-7^** | 5 | 0,54 |
| **HPC** | 67 | 12 | 6,49 | **1,31×10^-8^** | 8 | 0,22 |
| **mITC** | 73 | 13 | 5,02 | **3,60×10^-6^** | 6 | 0,27 |
| **lVC** | 54 | 8 | 4,95 | **7,97×10^-6^** | 8 | 0,39 |
| **lOFC** | 70 | 10 | 4,26 | **6,45×10^-5^** | 7 | 0,21 |
| **vmPFC** | 54 | 11 | 3,98 | **2,10×10^-4^** | 4 | 0,22 |
| Sdm | 4 | 2 | 5,11 | 0,01 | 1 | 0,20 |
| SPC | 9 | 4 | 4,40 | 2,28×10^-3^ | 3 | 0,56 |
| ACC | 18 | 7 | 4,21 | **5,85×10^-4^** | 2 | 0,17 |
| MTCr | 111 | 11 | 3,20 | 1,82×10^-3^ | 2 | 0,05 |
| Mv | 27 | 6 | 3,19 | 0,00 | 3 | 0,19 |
| SPCm | 6 | 2 | 3,15 | 0,03 | 2 | 0,33 |
| PCC | 19 | 8 | 3,11 | 6,02×10^-3^ | 3 | 0,21 |
| pINS | 66 | 16 | 3,08 | 0,00 | 5 | 0,08 |
| ITCr | 36 | 8 | 3,03 | 4,60×10^-3^ | 2 | 0,33 |
| IPCv | 63 | 10 | 2,92 | 4,93×10^-3^ | 2 | 0,08 |
| IPCd | 19 | 5 | 2,79 | 0,01 | 1 | 0,16 |
| PFrd | 14 | 5 | 2,79 | 0,02 | 1 | 0,07 |
| PMrv | 31 | 7 | 2,74 | 0,01 | 3 | 0,13 |
| Mdl | 25 | 5 | 2,70 | 0,01 | 2 | 0,32 |
| PFrvl | 15 | 6 | 2,62 | 0,02 | 2 | 0,40 |
| PFcdm | 13 | 5 | 2,59 | 0,02 | 2 | 0,31 |
| MTCc | 172 | 13 | 2,41 | 0,02 | 5 | 0,06 |
| Sv | 20 | 4 | 2,08 | 0,05 | 1 | 0,10 |
| PMdm | 5 | 2 | 1,59 | 0,19 | 1 | 0,20 |
| MCC | 6 | 4 | 1,39 | 0,22 | 0 | 0,00 |
| VCs | 18 | 6 | 1,33 | 0,20 | 3 | 0,22 |
| PFcdl | 25 | 9 | 1,03 | 0,31 | 1 | 0,04 |
| STCc | 105 | 15 | 1,02 | 0,31 | 1 | 0,01 |
| Sdl | 15 | 4 | 0,60 | 0,56 | 1 | 0,13 |
| PCm | 12 | 5 | -0,03 | 0,98 | 1 | 0,08 |
| PMdl | 15 | 5 | -0,38 | 0,71 | 1 | 0,07 |
| PFrm | 14 | 8 | -0,40 | 0,70 | 0 | 0,00 |
| Mdm | 2 | 1 | -0,79 | 0,57 | 0 | 0,00 |
| VCrm | 8 | 3 | -1,40 | 0,20 | 0 | 0,00 |
| STCr | 77 | 12 | -1,72 | 0,09 | 3 | 0,04 |
| Cu | 4 | 1 | -2,17 | 0,12 | 1 | 0,60 |
